# Accurate prediction of nucleic acid and protein-nucleic acid complexes using RoseTTAFoldNA

**DOI:** 10.1101/2022.09.09.507333

**Authors:** Minkyung Baek, Ryan McHugh, Ivan Anishchenko, David Baker, Frank DiMaio

## Abstract

Protein-nucleic acid complexes play critical roles in biology. Despite considerable recent advances in protein structure prediction, the prediction of the structures of protein-nucleic acid complexes without homology to known complexes is a largely unsolved problem. Here we extend the RoseTTAFold end-to-end deep learning approach to modeling of nucleic acid and protein-nucleic acid complexes. We develop a single trained network, RoseTTAFoldNA, that rapidly produces 3D structure models with confidence estimates for protein-DNA and protein-RNA complexes, and for RNA tertiary structures. In all three cases, confident predictions have considerably higher accuracy than current state of the art methods. RoseTTAFoldNA should be broadly useful for modeling the structure of naturally occurring protein-nucleic acid complexes, and for designing sequence specific RNA and DNA binding proteins.

---

Current approaches to protein-nucleic acid complex structure prediction involve building models of the protein and nucleic acid components separately and then building up complexes using computational docking calculations [1–3]. RNA structure prediction has generally proceeded by first predicting the secondary structure (Watson/Crick base pairing) and then assembling the secondary structure into a tertiary structure [4–7]. More recently, deep learning methods have been used for contact prediction to aid in RNA structure determination [8], and to select models from ensembles generated using other structure sampling approaches [9]. Despite this progress, the prediction of the structure of nucleic acids and protein-nucleic acid complexes has lagged considerably behind the prediction of protein structures from their amino acid sequences, which has been transformed by the high accuracy AlphaFold and RoseTTAFold methods.

AlphaFold and RoseTTAFold take as input one or more aligned protein sequences, and successively transform this information in parallel 1D, 2D and – in the case of RoseTTAFold – 3D tracks, ultimately outputting three dimensional protein structures. The 10s to 100s of millions of free parameters in these deep networks are learned by training on large sets of proteins of known structures from the PDB. Both AlphaFold2 and RoseTTAFold can generate accurate models of not only protein monomers but also protein complexes, modeling folding and binding by successive transformations over hundreds of iterations. Given the overall similarities between protein folding and RNA folding, and between protein-protein binding and protein-nucleic acid binding, we reasoned that the concepts and techniques underlying AlphaFold [10] and RoseTTAFold [11] could be extended to prediction of the structures of nucleic acids and protein-nucleic acid complexes from sequence information alone. We set out to generalize RoseTTAFold to model nucleic acids in addition to proteins, and to learn the many new parameters required for general protein-nucleic acid systems by training on the structures in the PDB. A major question at the outset was whether there were sufficient nucleic acid and protein-nucleic acid structures in the PDB to train an accurate and general model; key to the success of AlphaFold are the hundreds of thousands of protein structures in the PDB, but there are an order of magnitude fewer nucleic acid structures and complexes.

The architecture of RoseTTAFoldNA is illustrated in Figure 1. It is based on the three-track architecture of RoseTTAFold [11], which simultaneously refine three representations of a biomolecular system: sequence (1D), residue-pair distances (2D), and cartesian coordinates (3D). In addition to several modifications to improve performance [preprint], we extended all three tracks of the network to support nucleic acids in addition to proteins. The 1D track in RoseTTAFold has 22 tokens, corresponding to the 20 amino acids, a 21st “unknown” amino acid or gap token, and a 22nd mask token that enables protein design; to these, we added 10 additional tokens, corresponding to the 4 DNA nucleotides, the 4 RNA nucleotides, unknown DNA, and unknown RNA. The 2D track in RoseTTAFold builds up a representation of the interactions between all pairs of amino acids in a protein or protein assembly; we generalized the 2D track to model interactions between nucleic acid bases and between bases and amino acids. The 3D track in RoseTTAFold represents the position and orientation of each amino acid in a frame defined by three backbone atoms (N, CA, C), and up to four chi angles to build up the sidechain. For RoseTTAFoldNA, we extended this to include representations of each nucleotide using a coordinate frame describing the position and orientation of the phosphate group, and 10 torsion angles which enable building up of all the atoms in the nucleotide. RoseTTAFoldNA consists of 36 of these three-track layers, followed by four additional structure refinement layers, with a total of 67 million parameters.

**Figure 1.**
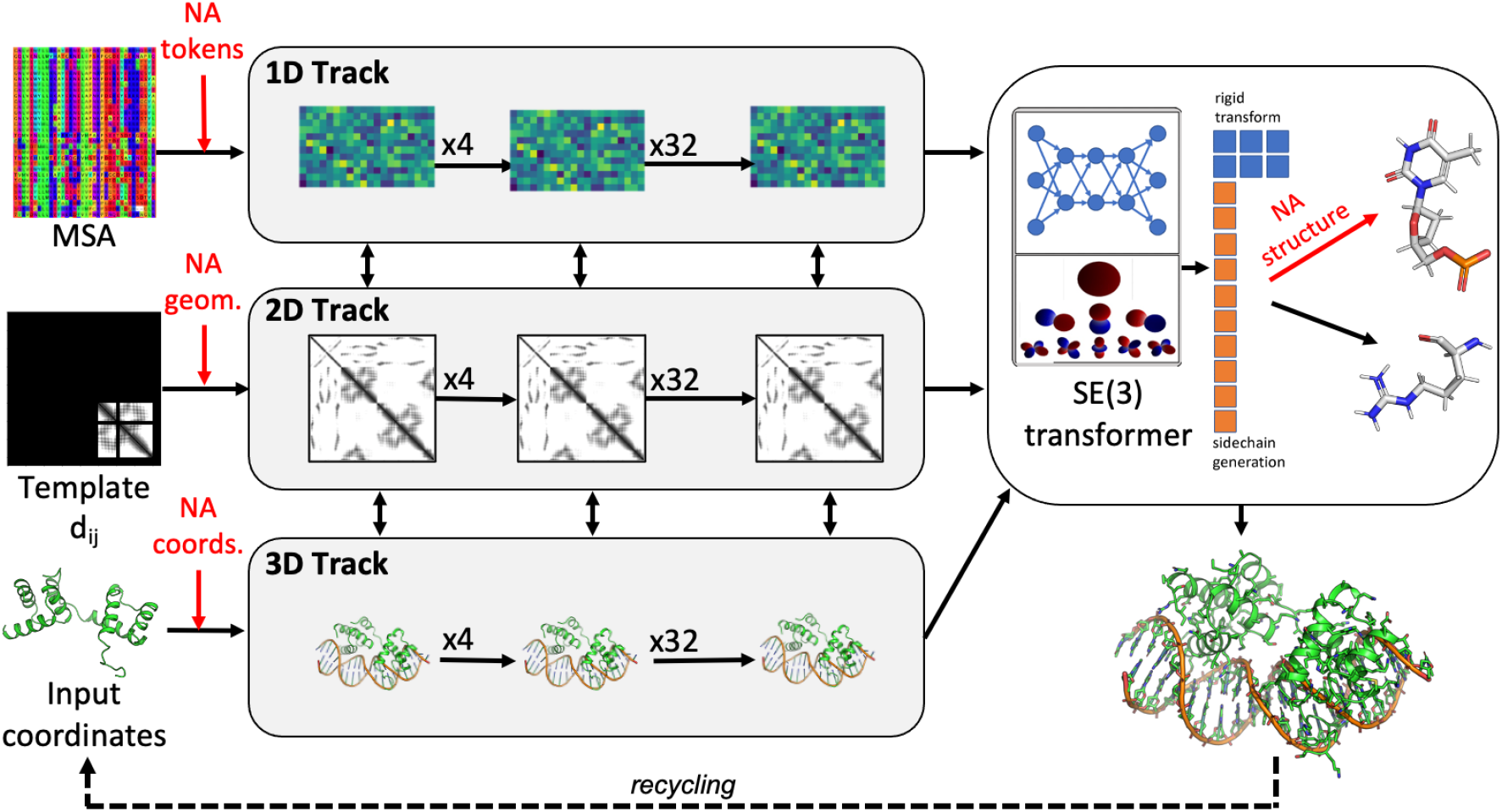
Overview of the architecture of RoseTTAFoldNA. The three-track architecture of RoseTTAFoldNA simultaneously updates sequence (1D), residue-pair (2D) and structural (3D) representations of protein/nucleic acid complexes. The areas in red highlight key changes necessary for the incorporation of nucleic acids: inputs to the 1D track include additional NA tokens, inputs to the 2D track represent template protein/NA and NA/NA distances (and orientations), and inputs to the 3D track represent template or recycled NA coordinates. Finally, the 3D track as well as the structure refinement module (upper right) can build all-atom nucleic acid models from a coordinate frame (representing the phosphate group) and a set of 10 torsion angles (6 backbone, 3 ribose ring, and 1 nucleoside).

We trained this end-to-end protein/NA structure prediction network using a combination of protein monomers, protein complexes, RNA monomers, RNA dimers, protein-RNA complexes, and protein-DNA complexes, with a 60/40 ratio of protein-only and NA-containing structures (see *Supplemental Methods*). Multichain assemblies other than the DNA double helix were broken into pairs of interacting chains. For each input structure or complex, sequence similarity searches were used to generate multiple sequence alignments (MSAs) of related protein and nucleic acid molecules. Network parameters were optimized by minimization of a loss function consisting of a generalization of the all atom FAPE loss [10] defined over all protein and nucleic acid atoms (see methods) together with additional contributions assessing recovery of masked sequence segments, residue-residue (both amino acids and nucleotides) interaction geometry, and error prediction accuracy. To try to compensate for the far smaller number of nucleic acid containing structures in the PDB (following sequence similarity based cluster to reduce redundancy, there are 1632 RNA clusters and 1556 protein-nucleic acid complex clusters compared to 26128 all protein clusters), we also incorporated physical information in the form of Lennard-Jones and hydrogen-bonding energies [12] as input features to the final refinement layers, and as part of the loss function during fine-tuning. During training, 10% of the clusters were withheld for model validation.

We trained the model using structures determined prior to May 2020, and used RNA and protein/NA structures solved since then as an additional independent validation set. For the validation set, complexes were not broken into interacting pairs and were processed entirely as full complexes. Paired MSAs were generated for complexes with multiple protein chains as described previously [13]. Due to GPU memory limitations, we excluded complexes with more than 1000 total amino acids and nucleotides, which resulted in 41 cases with a single RNA chain, 88 complexes with one protein molecule plus a single RNA chain (21) or DNA duplex (67), and 108 cases with more than one protein chain or more than a single RNA chain or DNA duplex.

The results of RoseTTAFoldNA at predicting RNA structures are summarized in Figure 2. Several observations are immediately apparent from these results. Most predictions are reasonably accurate: the average lDDT is 0.73, with 38% of models predicted with lDDT>0.8 (Figure 2A). RoseTTAFoldNA, like RoseTTAFold and AlphaFold, outputs not only a predicted structure but also a predicted model confidence, and as expected more confidently predicted models have higher accuracy: 42% of cases are predicted with very high confidence (plDDT > 0.9), for which the average lDDT is 0.84 and 72% of models have lDDT > 0.8 (and 96% lDDT > 0.7). Even for cases with no homologs of known structure or small numbers of sequence relatives (shallow MSAs), confidently predicted models are generally quite accurate (colorbar, Figure 2B,C). There were 78 cases with no detectable sequence similarity (>0.001 blastn E-value) to any structure in the training set, which have an average all-atom lDDT of 0.67, and 21% of models with lDDT>0.8. Within this set, 28% of cases are predicted with high confidence, with an average lDDT of 0.79 and 48% with lDDT > 0.8 (and 90% lDDT > 0.7). Four examples of high accuracy structure predictions in the absence of sequence homology to previously solved structures are shown in Figs 2D-G: these include a heavily modified bacterial ribosomal RNA, domain 2 of an NAD+ riboswitch, a lysine riboswitch, and the SARS-CoV-2 frameshifting pseudoknot, shown in Figures 2D-G. Inaccuracies in these predictions are in the curvature of larger models (Figures 2F,G) or near the binding site for the unmodeled ligand (Figures 2E), but the overall secondary structures and folds are correct.

**Figure 2.**
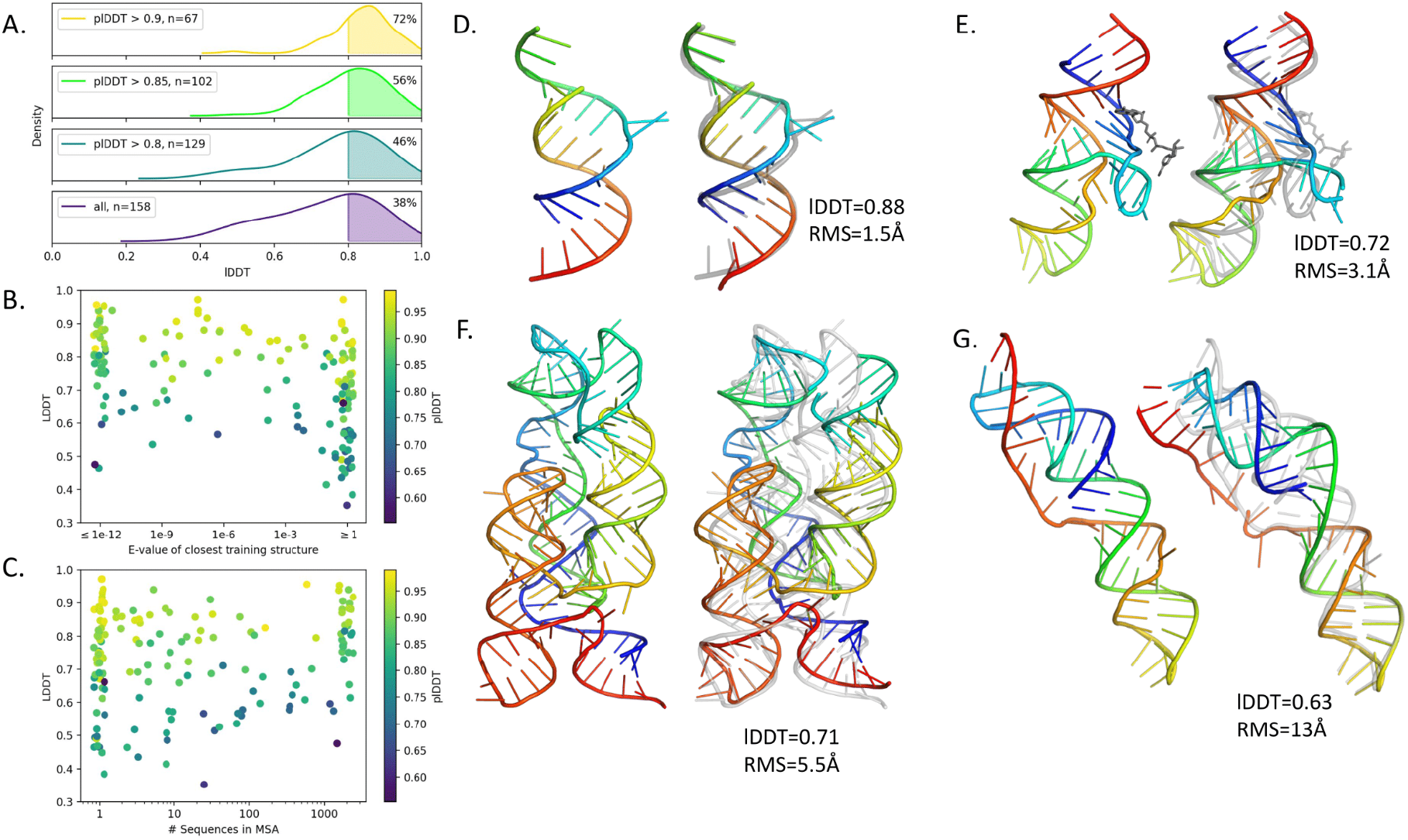
RNA structure prediction. (A-C) Summary of results on 117 RNA structures from the validation set and 41 RNA-only structures released since May 2020. (A) Model accuracy increases at higher confidence levels. The overall average lDDT is 0.73, and the average lDDT for very high confidence predictions (plDDT > 0.9) is 0.84. (B) Although models are generally more accurate for structures with close homologues in the training set, many structures with no detectable homologues in the training set are also predicted accurately. (C) Accuracy improves as the number of sequences in the MSA increases, but many single-sequence examples are accurately predicted. (D-F) Four example predictions of RNA models with no detectable sequence homologs in the training set, three of which also have no detectable structural homology according to PDB structure similarity search. (D) a heavily modified bacterial ribosomal RNA (PDB id: 2a04) [], (E) domain 2 of an NAD+ riboswitch (PDB id: 7d81) [], (F) a lysine riboswitch (PDB id: 3d0u) [], and (G) a SARS-CoV-2 frameshifting pseudoknot RNA (PDB id: 7lyj) [].

RoseTTAFoldNA results on 259 monomeric protein/NA complexes are summarized in Figure 3. As with the RNA cases, the predictions are reasonably accurate, with an average lDDT of 0.73 and 24% of models with lDDT>0.8 (Figure 3A), and about 56% of models identify greater than half of the native contacts between protein and NA (f_nat > 0.5, Figure 3C). As in the RNA case, and importantly for applications, the method correctly identifies which structure models are accurate. Although only 27% of the complexes are predicted with high confidence (mean interface PAE < 10), of those, 82% correctly model the protein/NA interface (“acceptable” or better by CAPRI metrics [14]). Over the 63 cases with no detectable sequence similarity to training protein/NA structures, the accuracy is somewhat lower on average (average lDDT=0.66 with 8% of models >0.8 lDDT and 31% with f_nat > 0.5), but the model is still able to correctly identify accurate predictions—all three of the high confidence predictions in this subset have acceptable interfaces according to CAPRI metrics. Four predictions of structures with no homology are shown in Figures 3D-G. These include a monomeric mutant of the transcriptional repressor BEND3, the endonuclease EndQ, the hMettl16 catalytic domain bound to an RNA hairpin, and Rmd9 bound to ssRNA. Inaccuracies in these predictions can be found in flexible linker regions (Figure 3D), distortions of the DNA double helix and a 1-base pair shift along the DNA (Figure 3E), translations of the interface (Figure 3F), and positions of flexible ssRNA termini (Figure 3G), but the overall complex architecture is clearly correctly recapitulated.

**Figure 3.**
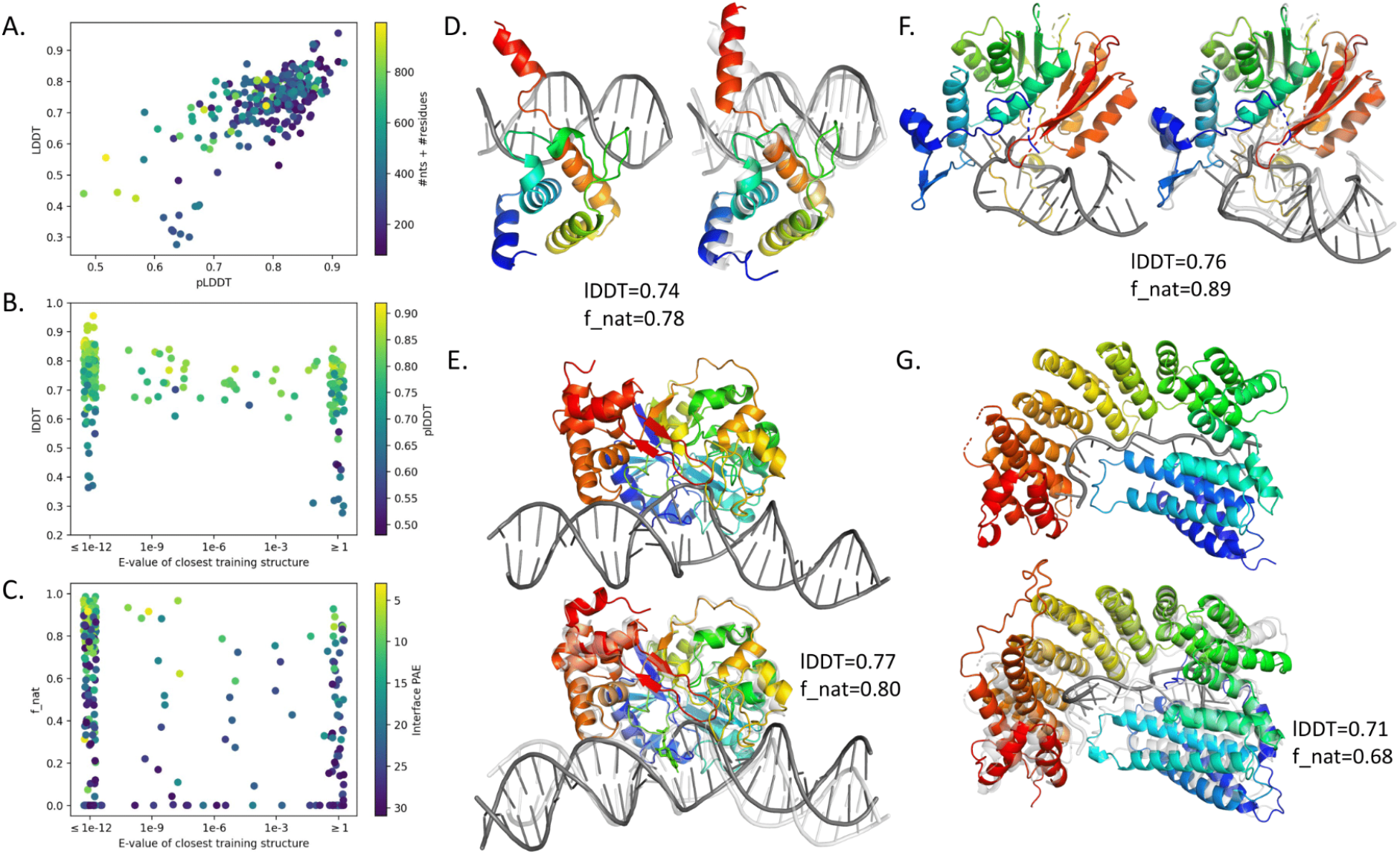
Protein - nucleic acid structure prediction. (A-C) Summary of results on 171 Protein/NA structures from the validation set and 88 Protein/NA structures released since May 2020. (A) Scatterplot of prediction accuracy (true lDDT to native structure) vs prediction confidence (lDDT predicted by the model) shows that the model correctly identifies inaccurate predictions. (B) Although accuracy is higher for models with close homologs, many models without any homologs are still predicted accurately. (C) Scatterplot of native interface contacts recapitulated in the prediction (f_nat) versus sequence similarity to training data. 31% of predictions are ranked “acceptable” or better by CAPRI metrics, and 82% of those with high confidence (mean interface PAE < 10). (D-G) Four examples of Protein/NA complexes without homologues: the transcriptional repressor BEND3 (panel D, pdb ID: 7v9i) []; the endonuclease EndQ (panel E, PDB id: 7k33) []; the hMettl16 catalytic domain bound to a 3’ UTR hairpin RNA (panel F, PDB id: 3du5); and Rmd9 – a protein that binds 3’ UTRs – bound to mRNA (panel G, PDB id: 7a9x).

**Figure 4.**
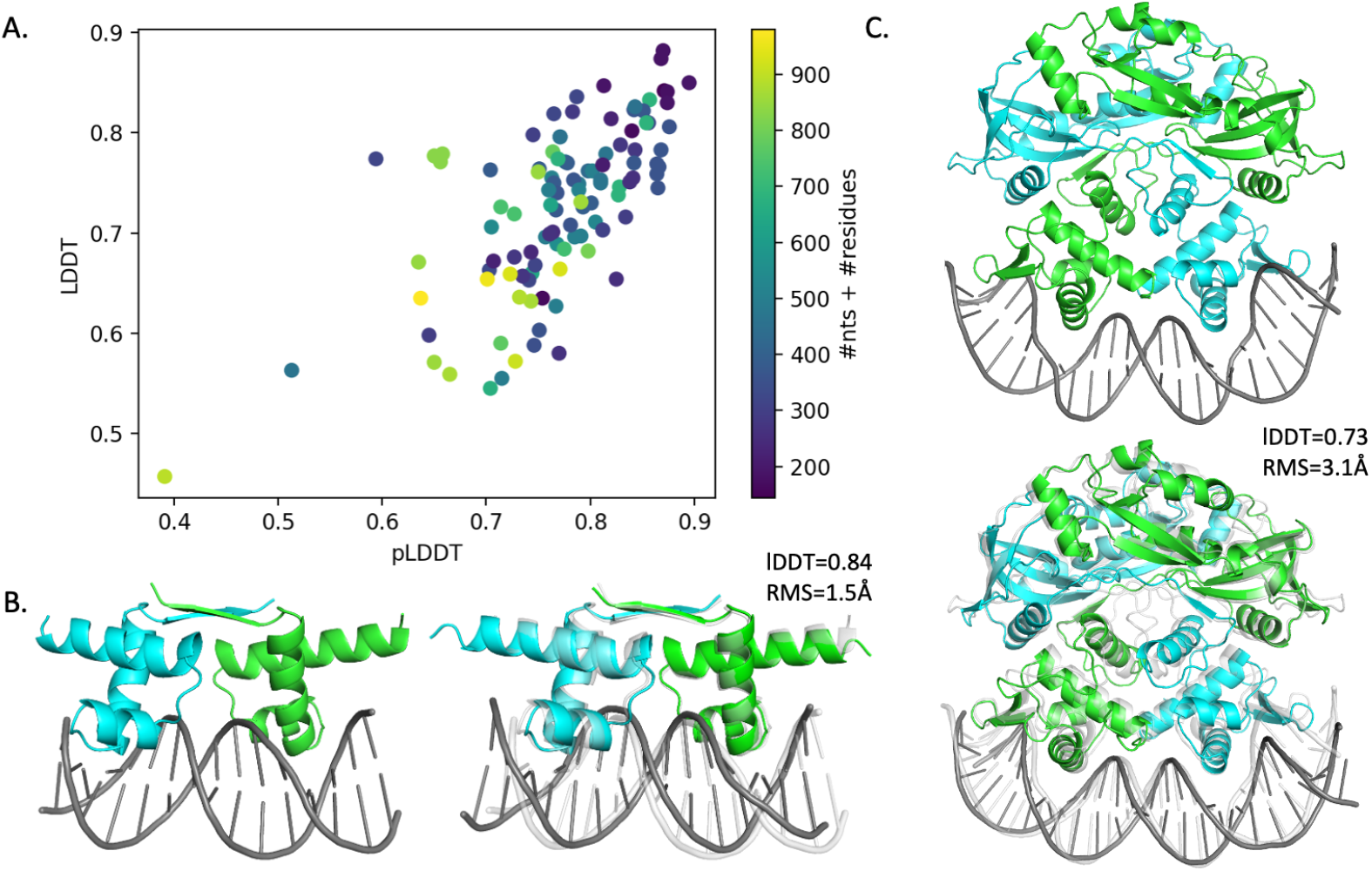
Modelling multichain protein-nucleic acid complexes. (A) Scatterplot of predicted model accuracy versus actual model accuracy for 108 protein/NA complexes with multiple protein chains or multiple nucleic acid chains/duplexes correctly estimates model error. (B-C) Examples of successful predictions without homologues in the training set. These include the dimeric bacterial transcription regulator DeoR (panel B, PDB id: 7bhy) []; and a dimeric complex from the BREX phage restriction system (panel C, PDB id: 7t8k) [].

Figure 5 summarizes the performance of RoseTTAFoldNA on 108 multi-subunit protein/NA complexes, most of which are either homodimers (66) or heterodimers (12) bound to nucleic acids. Performance is similar to that for monomeric protein/nucleic acid complexes, with an average lDDT=0.73 with 18% of cases >0.8 lDDT, and good agreement between confidence and accuracy (Figure 5A). Two examples are illustrated in Figure 5 (B,C), showing the ability of the model to predict complex structure as well as the “bending” of DNA induced by protein binding. Such bending would not be possible to predict by approaches that first generate models of the independent components and then rigidly dock them.

## Discussion

At the outset of this work, it was not clear that there were enough protein-nucleic acid structures in the PDB to enable robust training of a predictor with atomic accuracy – the training data used for nucleic acid prediction is only one tenth the size of the dataset used for protein structure prediction. Our results show, however, that this data is sufficient in many cases for de novo structure modeling, with RNA prediction accuracy nearly as good as protein structure prediction accuracy, and accurate modeling of protein/NA interfaces without shared MSA information or homologues of known structure in about 31% of cases. Prospective and blind tests will be important for further critical evaluation of the method.

Comparisons to the current state of the art methods are more difficult than was the case for the deep learning methods AlphaFold2 and RoseTTAFold which focused on the much more well studied protein structure prediction problem. There has been recent work on RNA structure prediction–over similar types of targets our method is much better than traditional methods and slightly better than other machine learning methods. Over the 78 RNA structures with no homologs in our training data, FARFAR2’s top-ranked models have an average lDDT of 0.47 [4], while DeepFoldRNA has an average lDDT of 0.61 [15], compared to 0.67 for RoseTTAFoldNA and 0.79 for RoseTTAFoldNA high-confidence predictions (Suppl. Fig S1; some of these examples may have been in the DeepFoldRNA training set). For protein-nucleic acid complexes, comparisons are even more difficult; indeed we are aware of no previous methods that predict the structures of such assemblies from sequence information alone, and there are not well established protocols for docking predicted protein and nucleic acid structures. Hence, while the accuracy of RoseTTAFoldNA on RNA structures and protein-nucleic acid complexes is considerably lower than that of AlphaFold2 on protein structures, it represents a very significant improvement in the state of the art.

Further increases in accuracy could come from a larger, more expressive network; we used a similar-sized network to that of RoseTTAFold2, with ^~^67M parameters and 36 total layers and there are more interactions to learn with the addition of nucleic acids. Use of high-confidence predicted structures as additional training examples (made more difficult by subsampling MSAs) should further increase model accuracy [10]; for this purpose there are databases of structured RNAs [15,16] and DNA binding profiles for thousands of proteins [17,18], and the latter should be useful for training a model fine-tuned for DNA specificity as well. Deep-learning guided structure prediction of proteins opened up new avenues of research; we hope that this work does the same for protein/NA interactions and complexes. To this end, we have made the method freely available at https://github.com/uw-ipd/RoseTTAFold2NA.

## Acknowledgements

This work was supported by Microsoft (M.B., D.B., and generous gifts of Azure computing time), the Audacious Project at the Institute for Protein Design (R.M., F.D., D.B.), National Science Foundation Grant #CHE 2226466 (F.D., D.B.), and the Howard Hughes Medical Institute (D.B.).

## Supplemental Methods

### Training and validation data processing

The protein and protein complex data used in training was identical to that used in training RoseTTAFold2. Additional data from RNA and protein/nucleic acid complexes was added to this. To construct this dataset, all PDBs solved by NMR, crystallography, or cryoEM at better than 4.5Å resolution were collected. A dataset was constructed considering all PDB structures published at or before April 30, 2020, and collecting:

· All RNA single chains and all RNA duplexes. A duplex was defined by looking for pairs of RNA chains making at least 10 hydrogen bonds.
· All interacting protein/nucleic acid pairs. Interacting pairs were defined by counting the number of 7Å contacts between protein Cαs and any (non-hydrogen) nucleic acid atom; if there were more than 16 such contacts, the pair was considered interacting. Nucleic acid duplexes were included if the DNA or RNA chains made at least 10 hydrogen bonds.

For modeling, the full-length sequence was used. All nonstandard bases/amino acids were converted into a backbone-only “unknown” residue type. The dataset size was 7396 RNA chains and 23583 complexes. These were then clustered using a 1e-3 hhblits [19] E-value for proteins and 80% sequence identity for RNA molecules, yielding 1632 nonredundant RNA clusters and 1556 nonredundant protein/NA clusters. These clusters were then split into training and validation sets, with clusters chosen for the training set; an example which contained any member (NA or protein) of a validation set cluster was assigned to the validation set. This led to 199 protein/NA clusters and 116 RNA clusters in the validation set.

Multiple sequence alignments (MSAs) were then created for all protein and RNA sequences in the training and validation set. Protein MSAs were generated in the same way as RoseTTAFold [preprint], using hhblits at successive E-value cutoffs (1e-30, 1e-10, 1e-6, 1e-3), stopping when the MSA contains more than 10000 unique sequences with >50% coverage. RNA MSAs were generated using a pared-down version of rMSA (https://github.com/pylelab/rMSA) that removes secondary structure predictions: sequences were searched using blastn [20] over 3 databases (RNAcentral [15], rfam [16], and nt) to first identify hits, then using nhmmer [21] to rerank hits. We again use successive E-value cutoffs (1e-8, 1e-7, 1e-6, 1e-3, 1e-2, 1e-1), stopping when the MSA contains more than 10000 unique sequences with >50% coverage.

### Test set data processing

For an independent test set, we took all structures published to the PDB May 1, 2020 or later. Selection criteria and preprocessing was the same as for the training and validation data with two exceptions: a) only complexes fewer than 1000 residues plus nucleotides in length were considered; and b) for complexes containing more than one unique protein chains, paired MSAs were created by merging sequences from the same organism into a single combined sequence (following prior work [13]). This gave us 93 complexes with one protein molecule plus a single RNA chain or DNA duplex, 41 cases with a single RNA chain, and 132 cases with more than one protein chain or more than a single RNA chain or DNA duplex.

### Loss functions

The model was trained using a loss function similar to RoseTTAFold, where we take the weighted sum:

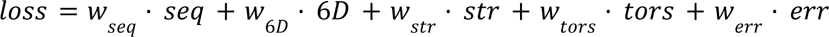

Above, seq is the masked amino-acid recovery loss (no masking is applied to nucleotide sequences); 6D is the 6-dimensional “distogram” loss [ref - trRosetta]; str is the structure loss, consisting of the average backbone FAPE loss [10] over all 40 structure layers of the network plus the allatom FAPE loss for the final model; tors is the torsion prediction loss averaged over the 40 structure layers; and err is the loss in pLDDT prediction.

FAPE loss is extended to nucleic acids in a straightforward manner from how it is implemented for amino acids. For backbone FAPE loss, the phosphate group in the nucleic acid backbone is treated as the nucleotides “frame,” in the same way that N-Cα-C is used as an amino acid frame. For nucleic acid allatom FAPE loss, three-atom frames are constructed corresponding to each of the 10 “rotatable torsions” (see below for the definition), where the frame consists of the two bonded atoms defining the torsion plus an additional bonded atom, closer to the phosphate group in the bond graph. The cross product of these 10 frames with all atoms is used to calculate FAPE loss.

Following training with the above loss function, an additional “finetuning” phase is carried out, where additional energy terms are added to the loss function enforcing reasonable model geometry:

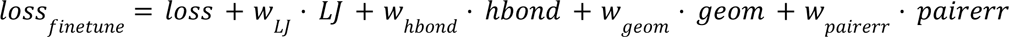

Above, LJ and hbond are the Lennard-Jones and hydrogen bond energies of the final structure (normalized by the number of atoms), using a reimplementation of the corresponding Rosetta energy terms [12]; geom is a term that enforces ideal bond lengths and bond angles around the peptide or phosphodiester bond connecting residues/nucleotides; and pairerr is a predicted residue-pair error [10]. The functional form of the geom term is identical to that of RoseTTAFold2, a linear penalty with a “flat bottom” plus or minus 3 degrees/0.02 Å from the ideal values.

### Model training

The network was trained in two stages, an initial training period, and a fine-tuning period. In both, input structures were divided into 5 pools: a) protein structures, b) “distilled” protein structures (consisting of high-confidence AlphaFold2 predictions), c) protein complexes, d) protein/NA complexes, and e) RNA structures. Training sampled from each of these pools with equal probability (though later in training protein/NA frequency was increased to 25% and RNA frequency lowered to 15%). For both pools containing “complexes,” an equal number of positive and negative examples were used in training. Negative examples consist of non-binding proteins or protein/NA pairs; the structure loss only penalizes each component individually, and the 6D loss favors placing negative binding examples far apart.

Examples larger than 256 residues/nucleotides in length were “cropped” to 256 residues in length. For protein-only data these crops were continuous sequences; for nucleic acids and nucleic-acid/protein complexes the cropping was a bit more complex. A graph was constructed where sequential residues/nucleotides had edges with weight 1, Watson/Crick base-paired nucleotides had weight 0, and protein/NA bases closer than 12Å (Cα to P) had a weight of 0. In negative cases, a single random protein/NA edge was given weight 0. Then minimum-weight graph traversal starting from a randomly chosen protein/NA edge was used to crop the model down to 256 residues/nucleotides. For RNA-only models the same strategy was used, though the starting point was a random nucleotide.

Training was carried out in parallel on 64 GPUs. A batch size of 64 was used throughout training with a learning rate of 0.001, decaying every 5000 steps. The following weights were used: w_seq_=3.0, w_6d_=1.0, w_str_=10.0, w_tors_=10.0, w_err_=0.1. The Adam optimizer was used, with L2 regularization (coeff=0.01).

Following ^~^1e5 optimization steps, fine-tuning training was carried out. Here we increase crop size to 384 and effective batch size to 128, and reduce learning rate to 5e-4. We used additional loss terms with weights w_geom_=0.1, w_LJ_=0.02, w_hbond_=0.05, and w_pairerr_=0.1, and optimized for an additional 30000 minimization steps. All told, training took approximately 4 weeks.

---

**Supplemental Figure S1.**
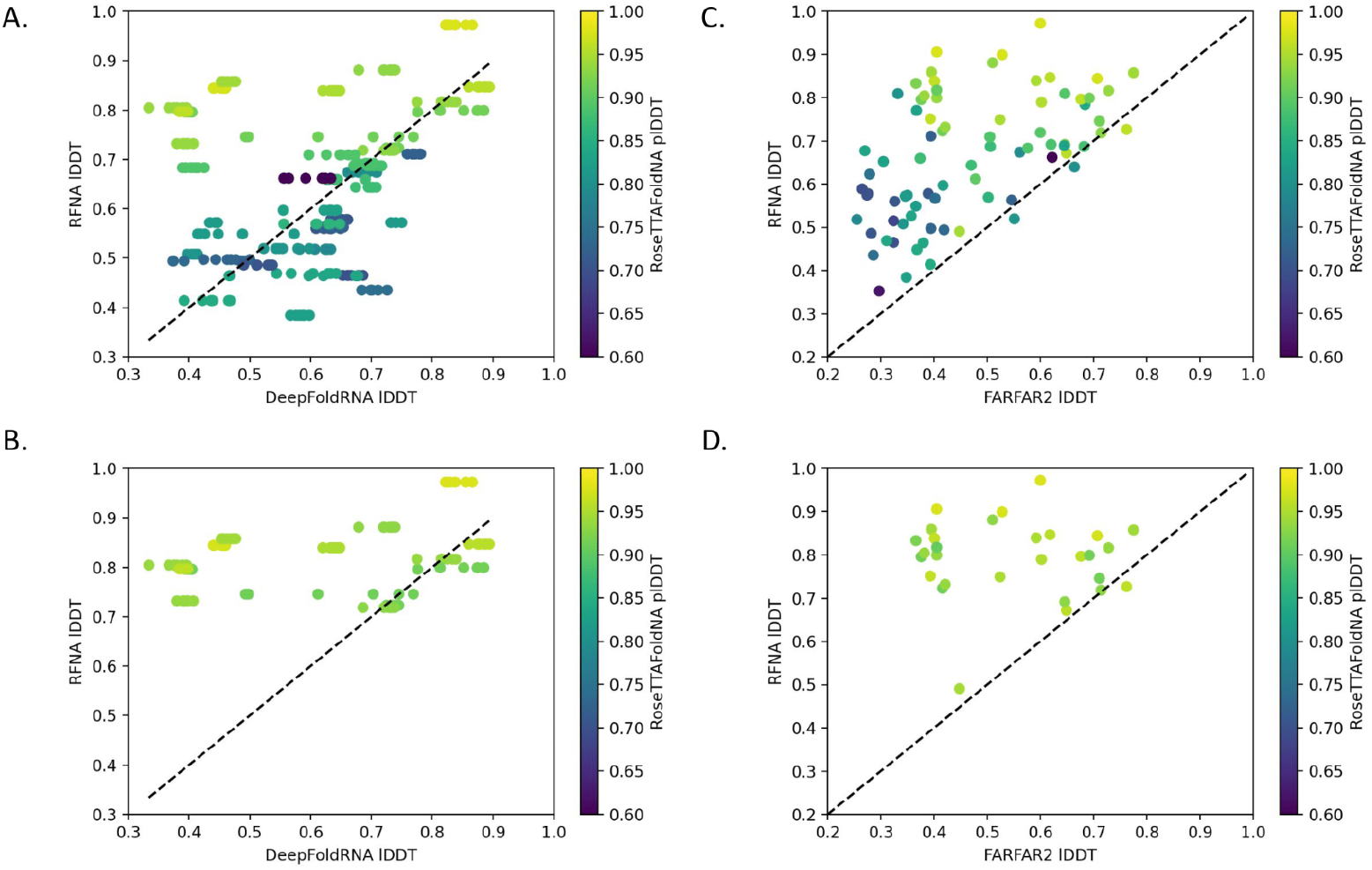
Comparing RoseTTAFoldNA to other methods. (A) Scatterplot of predicted accuracy for RoseTTAFoldNA vs DeepFoldRNA, a recent machine learning method for RNA structure prediction [15] shows comparable performance. RoseTTAFoldNA slightly outperforms DeepFoldRNA, with average lDDTs of 0.67 and 0.61 respectively. (B) The gap in performance is much larger if only RoseTTAFold’s high-confidence predictions (plDDT > 0.9?) are considered, which have an average lDDT of 0.79. (C) Scatterplot comparing RoseTTAFoldNA to FARFAR2, a Rosetta-based fragment assembly method for RNA structure prediction [4]. RoseTTAFoldNA consistently and dramatically outperforms FARFAR2’s top-ranked models, which have an average lDDT of 0.47. (D) The performance gap is even larger when only considering RoseTTAFoldNA confident predictions.

